# New insights on human essential genes based on integrated analysis

**DOI:** 10.1101/260224

**Authors:** Hebing Chen, Zhuo Zhang, Shuai Jiang, Ruijiang Li, Wanying Li, Chenghui Zhao, Hao Hong, Xin Huang, Hao Li, Xiaochen Bo

**Author notes:** To whom correspondence should be addressed; (H.L.); (X.B). These authors contributed equally to this work (Hebing Chen, Zhuo Zhang, Shuai Jiang, Ruijiang Li).

## Abstract

Essential genes are those whose functions govern critical processes that sustain life in the organism. Recent gene-editing technologies have provided new opportunities to characterize essential genes. Here, we present an integrated analysis for comprehensively and systematically elucidating the genetic and regulatory characteristics of human essential genes. First, essential genes act as “hubs” in protein-protein interactions networks, in chromatin structure, and in epigenetic modifications, thus are essential for cell growth. Second, essential genes represent the conserved biological processes across species although gene essentiality changes itself. Third, essential genes are import for cell development due to its discriminate transcription activity in both embryo development and oncogenesis. In addition, we develop an interactive web server, the Human Essential Genes Interactive Analysis Platform (HEGIAP) (http://sysomics.com/HEGIAP/), which integrates abundant analytical tools to give a global, multidimensional interpretation of gene essentiality. Our study provides a new view for understanding human essential genes.

## Introduction

Essential genes are indispensable for organism survival and maintaining basic functions of cells or tissues (1–3). Systematic identification of essential genes in different organisms (4) has provided critical insights into the molecular bases of many biological processes (5). Such information may be useful for applications in areas such as synthetic biology (6) and drug target identification (7, 8). The identification of human essential genes is a particularly attractive area of research because of the potential for medical applications (9, 10). Utilizing gene-editing technologies based on CRISPR-Cas9 and retroviral gene-trap screens, three independent genome-wide studies (11–13) identified essential genes that are indispensable for human cell viability. The results agreed very well among the three studies, confirming the robustness of the evaluating approaches. All of the studies (11–13) found that ~10% of the ~20,000 genes in human cells are essential for cell survival, highlighting the fact that eukaryotic genomes have intrinsic mechanisms for buffering against genetic and environmental insults (14).

In a recent review, Norman Pavelka and colleagues (15) stated that gene essentiality is not a fixed property but instead strongly depends on the environmental and genetic contexts and can be altered during short- or long-term evolution. However, this paradigm leaves some confusion, as we do not know how essential genes are interconnected within the cell, why these genes are essential, or what their underlying mechanisms may be. Furthermore, we do not know whether these genes are associated with disease (e.g., cancer) or have the potential to be exploited as targets for therapeutic strategies. With the recent and rapid development of next-generation sequencing and other experimental technologies, we now have access to myriad data on genomic sequences, epigenetic modifications, structures, and disease-related information. These data will enable researchers to examine human essential genes from multiple perspectives.

In light of this background, we performed a comprehensive study of human essential genes, including their genomic, epigenetic, proteomic, evolutionary, and embryonic patterning characteristics. Genetic and regulatory characteristics were studied to understand what makes these genes essential for cell survival. We analyzed the evolutionary status of human essential genes and their profiles during embryonic development. Our findings suggest that human essential genes are important for lineage segregation. Essential genes have important implications for drug discovery, which may inform the next generation of cancer therapeutics (Fig. 1). Finally, we have developed a new web server, the Human Essential Genes Interactive Analysis Platform (HEGIAP), to facilitate the global research community in the comprehensive exploration of human essential genes.

**Figure 1.**
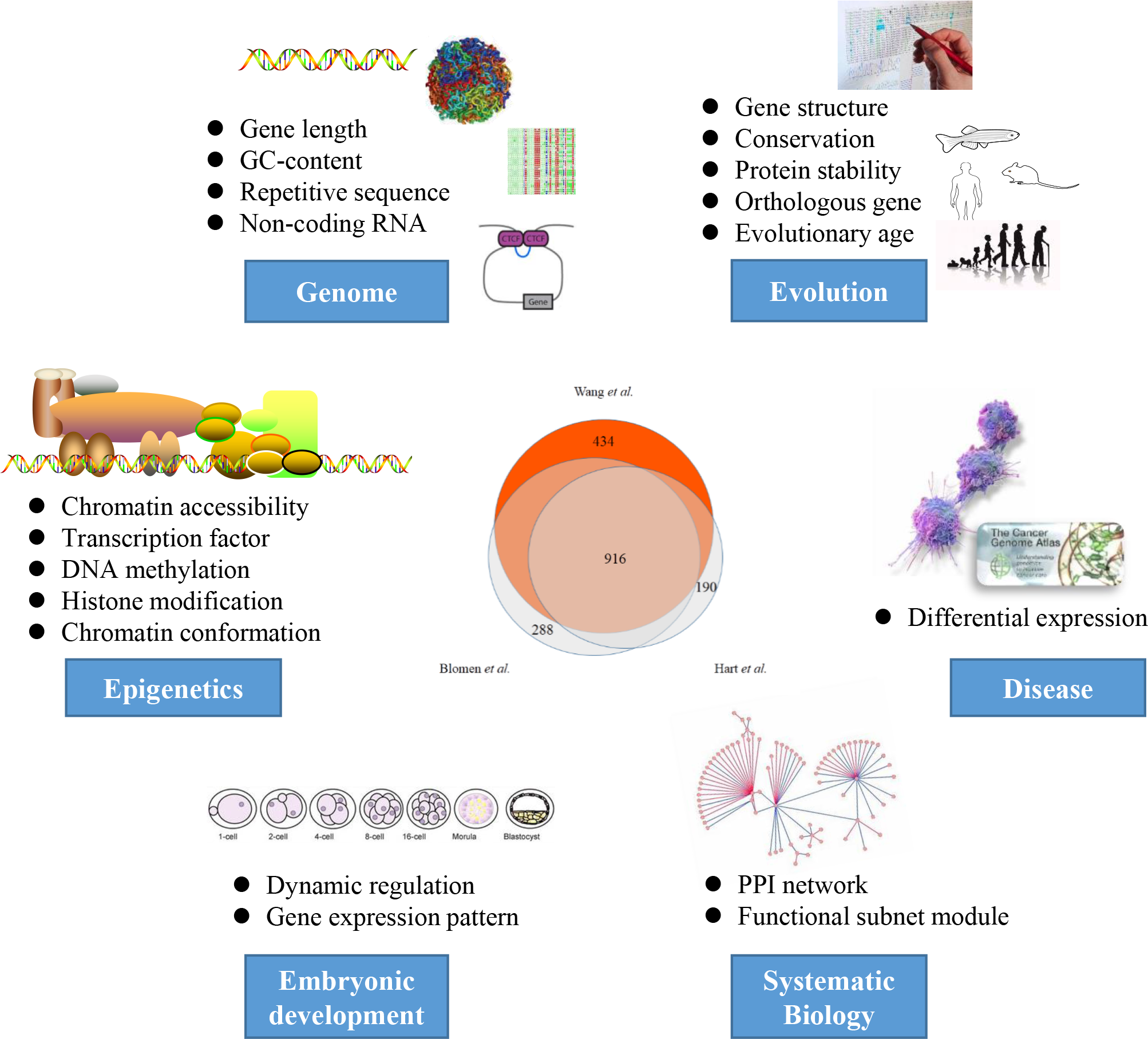
Comprehensive overview of the integrated analysis of human essential genes

## Results

### Multi-level essentiality of human essential genes

Essential genes define the key biological functions that are required for cell growth, proliferation, and survival. To characterize human essential genes, we first compared three essential gene sets generated by different experimental methods (11–13), over 60% of essential genes were cataloged in at least two literatures (Venn diagram in Fig. 1). Here, we focused on the essential genes detected by Wang et al (11), as it contains more than two-thirds of the essential genes in the other two studies, and defines a CRISPR score (CS) to assess the gene essentiality, where low CS indicates high essentiality. The sets of essential genes in multiple cell lines showed a high degree of overlap (71.78%, 77.26% and 72.36% of essential genes in KBM7 cell line are also essential in other 3 cell lines respectively), here we used the data from the KBM7 cell line for further analysis. We then divided all protein-coding genes into ten groups (from CS0 to CS9) according to the ascending CS values, where group CS0 indicates essential genes. This detailed classification provided a well representation of the various features in subsequent analyses.

#### Protein essentiality: High transcription activation and stability, “hub” of protein-protein interaction network

We first analyzed the expression level of human genes in 2,916 individuals from the GTEx program (16). Essential genes were highly expressed compared to nonessential genes (*p*-value < 1×10^−50^, Welch’s t-test) and the expression level decreased as gene essentiality decreased (the CS value increased) (R^2^ = 0.42, *p*-value = 0.04) (Fig. 2A). Furthermore, proteins encoded by essential genes showed higher stability compared to other proteins (*p*-value = 1.10×10^−18^, Welch’s t-test) (fig. S1A). This observation is consistent with a recent study (17), which found that highly expressed proteins are stable because they are designed to tolerate translational errors that would lead to accumulation of toxic misfolded species.

**Figure 2.**
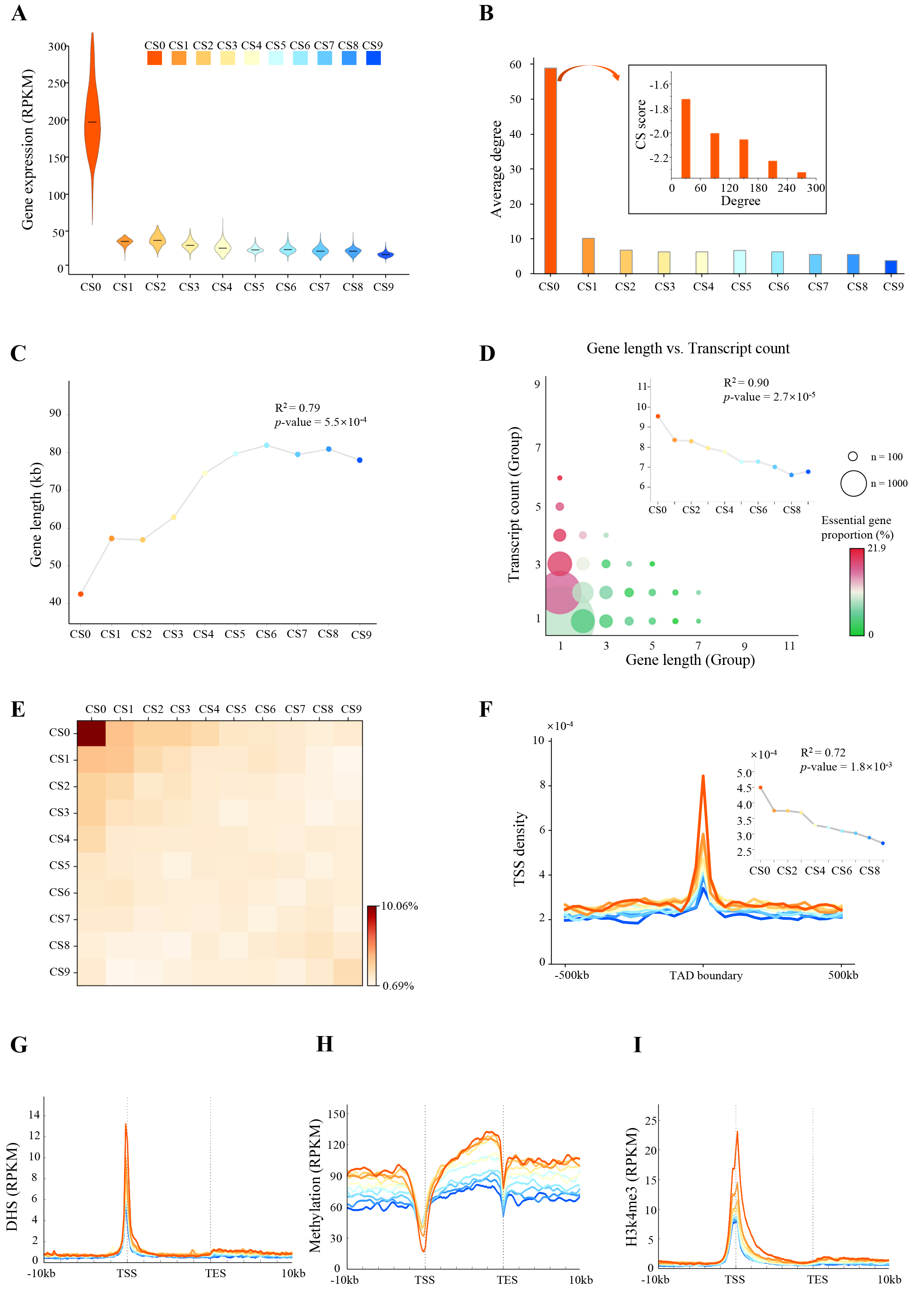
General properties of human essential genes. (A) Violin plot showing gene expression of 2916 individuals from GTEx program. Mean expression level was calculated for each group of genes (CS0-CS9). (B) Degree of connectivity for each gene group. Inset: relationship between degree of connectivity and CS value of group CS0 (human essential genes). (C) Relationship between gene essentiality and gene length. R^2^ and *p*-value for linear regression are shown. (D) Relationship between gene length and transcript count. R^2^ and *p*-value for linear regression are shown. (E) Heatmap showing the colocalization of gene’s TSS. (F)TSS density surrounding TAD boundary (IMR90 cell line). Inset: average TSS density within regions: 50 kb upstream, TAD boundary and 50 kb downstream. (G-I) Profiles showing mean signal for chromatin accessibility **(G)**, methylation level (**H**), and H3k4me3 density **(I)**.

In many model organisms, essential genes tend to encode abundant proteins that engage extensively in protein-protein interactions (PPIs) (18). We constructed a PPI network for each group (CS0-CS9) (fig. S1B), the connectivity degree of essential genes were significant higher than other genes (*p*-value = 6.50×10^−178^, Welch’s t-test) (Fig. 2B) and the degree was negatively correlated with CS value in essential gene set (R = −0.23, *p*-value = 9.70×10^−25^) (insets in Fig. 2B). We then calculated the distribution of genes with ranged connectivity degree. We found that, with higher degree, there are less genes (fig. S2A) but higher proportion of essential genes (fig. S2B). These results indicated that essential genes located at central hubs of the PPI networks. We finally performed a Gene Ontology (GO) analysis for each group, essential genes were enriched in fundamental biological processes, such as rRNA processing, translational initiation, mRNA splicing, and DNA replication while nonessential genes were less significant enriched in other processes (fig. S3).

In summary, essential genes are highly expressed and associated with important biological processes, proteins encoded by essential genes are stable and locate at connect hubs in PPI networks. These results together show the essentiality of essential genes at the protein level.

#### Structural essentiality: The high density in the genome and 3D structure lead to a “hub” of chromatin organization

In general, gene length affects the stability of the kinetics of genetic switches and thus affects the dynamics of gene expression (19), we found that human essential genes were much shorter than nonessential genes (*p*-value = 1.38×10^−53^, Welch’s t-test) (Fig. 2C), which is consistent with a previous study in *E. coli* (19). Generally, long genes are likely to contain abundant transcripts in human genome (R = 0.34, *p*-value = 1.0×10^−100^, Pearson correlation) (fig. S4A), however, more transcripts were found in essential genes compared to nonessential genes (*p*-value = 7.02×10^−42^, Welch’s t-test) (Fig. 2D) and transcript counts decreased as essentiality decreased (R^2^ = 0.90, *p*-value = 2.70×10^−5^, Pearson correlation) (insets in Fig. 2D), indicating that mRNAs transcribed by essential genes are highly variable. As reported, GC content is associated with DNA stability and variations in the GC ratio within the genome result in variations in staining intensity in chromosomes (18). We then examined the distribution of GC content. As expected, essential genes contained the highest GC content (fig. S4B). Further, repetitive elements in the genome have multiple functions, including a major architectonic role in higher-order physical structuring (20) and repetitive component of the genome can detect and repair errors and damage to the genome (21). We then investigated repetitive elements within essential genes. DNA sequences for interspersed repeats and low-complexity DNA sequences were first identified and masked by RepeatMasker (22). We found that repetitive elements were significant enriched in essential genes compared to that in nonessential genes (*p*-value < 1×10^−50^, Welch’s t-test) (fig. S4C). Together, essential genes are formed with high GC content and colocalize with repetitive elements, suggesting that essential genes are much stable and may help maintain genomic stability.

We next examined the genomic distribution of essential genes, the transcription start sites (TSSs) of essential genes are prone to be clustered (Fig. 2E, fig. S5A). We then investigated the three-dimensional (3D) structural organization of essential genes. Previous studies reported that topologically associated domains (TADs) are highly conserved between cell types and species and proximity to the TAD boundary likely contributes to the stabilization of gene expression (23–27). We detected TADs using high-throughput/high-resolution chromosome conformation capture (Hi-C) data from Ren (28) and Rao (29). We observed significant clusters of essential genes within TAD boundary compared to that of nonessential genes (*p*-value < 1×10^−16^, Welch’s t-test) (Fig. 2F, fig. S5B-E).

Chromatin may be associated with proteins having affinity for each other, resulting in chromatin loops (30). We then calculated the intra-TAD local (<100 kb) Hi-C contacts for each gene set and examined the distribution of gene’s TSSs located in chromatin loop anchors. Essential genes contained more local contacts (*p*-value < 1×10^−43^, Welch’s t-test) (fig. S6A-B) and TSSs density of essential genes that located in chromatin loop anchors was higher than that of nonessential gene groups (fig. S6C). As the formation of architectural loops is strongly dependent on the protein CTCF (31), we examined the distribution of CTCF signal detected by ChIP-seq (32). As expected, CTCF was more likely to bind near essential genes (fig. S6D).

In summary, essential genes are structural essential due to their high GC content, highly enriched repetitive elements and central location in the chromosomal scaffold.

#### Epigenetic essentiality: The enrichment of epigenetic marks lead to a “hub” of epigenetic regulatory network

Epigenetic modification of chromatin provides the necessary plasticity for cells to respond to environmental and positional cues, and enables the maintenance of acquired information without changing the DNA sequence (33). To study the epigenetic information on essential genes, we took advantage of the recent high-throughput genomic assays (34, 35). We first examined the chromatin accessibility of essential genes. Strong DHS signals were observed in the promoter of essential genes and such enrichment decreased as gene essentiality decreased (Fig. 2G). Moreover, more transcription factor binding sites (TFBSs) were detected in the promoter of essential genes (fig. S7). We then examined the DNA methylation pattern of essential genes. By analyzing data from two sequencing-based methods, DNA immunoprecipitation (MeDIP-seq) and methylation-sensitive restriction enzyme (MRE-seq) (36), we found that the methylation level of gene’s promoter increased while gene essentiality decreased (R^2^ = 0.93, *p*-value = 8.6×10^−6^ for MRE-seq, R^2^ = 0.93, *p*-value = 5.8×10^−6^ for MeDIP-seq, Welch’s t-test) (Fig. 2H and fig. S8A). Next, we examined two histone modifications including trimethylation of H3 lysine 4 (H3K4me3) and trimethylation of H3 lysine 27 (H3K27me3), which is associated with transcription activation and gene repression (37–39), respectively. Similar to chromatin accessibility, the H3K4me3 signals were strongly enriched in the promoters of essential genes and the H3K4me3 density of gene’s promoter increased while gene essentiality increased (Fig. 2I). In contrast, the H3K27me3 signals were weakest in the promoter of essential genes compared to nonessential genes (fig. S8B). Finally, we studied the abundance of noncoding RNAs (ncRNAs), which play a key role in regulating gene expression (40, 41). The density of ncRNAs in the promoter of genes decreased as essentiality decreased (R^2^ = 0.40, *p*-value = 0.049) (fig. S8C).

In summary, these results have provided epigenetic evidence demonstrating that essential genes are hubs of active epigenetic modifications.

### Evolutionary nature of human essential genes

Genes that express highly and widely are reported to be originated early and conserved across species (42). We next investigated the universal distribution of evolutionary rates of essential genes. Using gene origination times inferred in a recent research (21), we showed that, as expected, essential genes, on average, were older (Fig. 3A) and significant conserved (*p*-value = 9.40×10^−9^, Welch’s t-test) (fig. S9) than nonessential genes. However, a small subset of essential genes were notably young (Fig. 3B), we found that the proportion of essential genes in human-specific genes (gene age group 13) is significantly larger compared to that of other genes (gene age groups 1-11) (Fig. 3C), which means that human-specific genes are more likely to be essential genes in human. Similar results were also observed in other 3 cell types (fig. S10). GO analysis of essential genes in the youngest 2 gene groups (human and chimpanzee) showed a significant enrichment in the regulation of GTPase activity (fig. s11A). We also found that these youngest essential genes are shorter, have a lower conservation level and not as actively expressed as other essential genes (fig. S11B). In addition, “old” genes are reported to be longer than “young” genes (43–46), however, essential genes are mostly “old” genes but are shorter than other genes on average (Fig. 3D).

**Figure 3.**
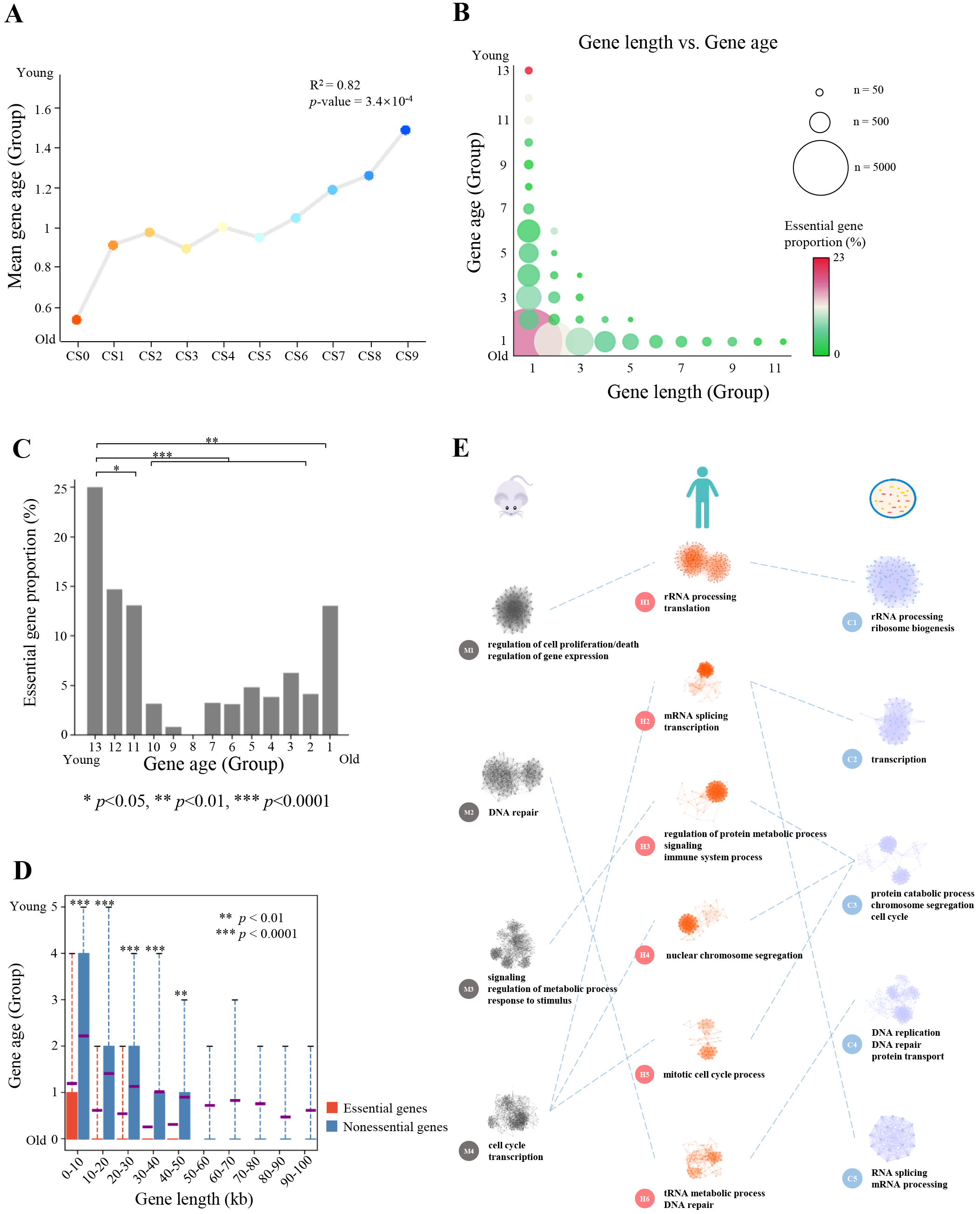
Evolutionary nature of human essential genes. (A) Relationship between gene essentiality and gene age. R^2^ and *p*-value for linear regression are shown. (B) Scatter-plot of gene length and evolutionary age. Size of the circle indicates number of genes; color represents essential gene proportion. (C) Essential gene proportion in all 13 gene age groups. (D) Analysis of gene age difference between essential genes and nonessential genes grouped by gene length. Red boxes: gene age data of essential genes. Blue boxes: gene age data of nonessential genes. Violet line: mean value. Gene group containing less than 100 genes are not shown. (E) Essential gene associated (EG-associated) specific functional modules. The network of specific functional interactions among the 1,878 human essential genes was clustered by using a graph theory clustering algorithm to elucidate gene modules. Six clusters that containing ≥40 genes (H1-H6) were tested for functional enrichment by using genes annotated with GO biological process terms. Representative processes and pathways enriched within each cluster are presented alongside the cluster label. Enriched functions provide a landscape of the potential effects of cellular functions for essential genes. Similar functional processes were shared by essential genes in mouse (four subnet modules) and *S. cerevisiae* (five subnet modules).

Evolutionary age was defined based on the presence of a homolog in a wide range of species from single-celled organisms to primates (47). However, the essentiality of a gene can change in the course of evolution (48). We further investigated the evolvable nature of gene essentiality in human and other 4 species including mouse, *Danio rerio*, *Drosophila melanogaster*, and *Saccharomyces cerevisiae*. To our surprise, roughly more than 80% of genes found to be essential in human were non-essential in other species and vice versa, similar results were observed in another human gene sets containing 8,253 essential genes(4) (fig. S12).

One possibility for the changes of essential genes in different species is that genes or functions could arise separately or be lost or replaced by others during evolution and thus the biological network could become more robust (48). To test this hypothesis, we compared PPI networks between species. A PPI network in each species was first constructed with essential genes using network topology method described previously (49) and subnet modules, namely the densely connected regions which can represent molecular complexes, were detected using MCODE algorithm (50). We then performed gene set enrichment analysis (GSEA) (51) for each subnet modules. Interestingly, similar biological processes were observed between human and other species although the essential genes were quite different (Fig. 3E), for instance, rRNA processing was enriched in human, mouse and *S. cerevisiae* but only less than 18% of essential genes in human were essential in mouse or *S. cerevisiae* (fig. S12). Our observations suggest that essentialomes are enriched in genes required for essential processes.

### Transcription activation of essential genes during cell development

#### Dynamic expression of essential genes in early embryo development

Cell fate decisions fundamentally contribute to the development and homeostasis of complex tissue structures in multicellular organisms. The key to answer how and why apparently identical cells have different fates lies in the emergence of transcriptional programs (52). We next characterized essential genes in mammalian embryonic development. Due to the lack of experimental data on human embryos, we used data from mouse preimplantation embryos. To examine whether the genetic and epigenetic features were consistent in human and mouse, we calculated the distribution of gene expression and epigenetic information in human and mouse embryonic development, strong correlation was observed between human and mouse (fig. S13). Thus, it is reasonable to use mouse data to study human essential genes. During embryos development, the expression level of essential genes were progressively increased and two significant increasing were observed in 2-cell embryos and inner cell mass (ICM), which corresponding to zygotic genome activation (ZGA) (53, 54) and the first cell fate decision (55), respectively. In contrast, the expression level of nonessential genes were significant lower than essential genes during the entire preimplantation (*p*-value < 10^−100^, Welch’s t-test) and genes in groups CS7-CS9, which labeled as the least essentiality, were silent after 2-cell embryos, which similar to the maternal mRNA degradation process (Fig. 4A). To further understand the dynamic changes of transcription activity of essential genes during embryos development, we investigated the dynamics of chromatin state. The accessible chromatin and active histone modifications were highly enriched in the promoters of essential genes compared to nonessential genes (Fig. 2G, I and Fig. 4B, C). In addition, chromatin was progressively accessible and the H3K4me3 density was progressively increased (Fig. 4B, C). However, essential genes were least methylated during embryos preimplantation (Fig. 4E). These observations suggest that essential genes are required to be expressed for embryo development and both chromatin accessibility and epigenetic modifications may contribute to the formation of transcriptional programs in essential genes.

**Figure 4.**
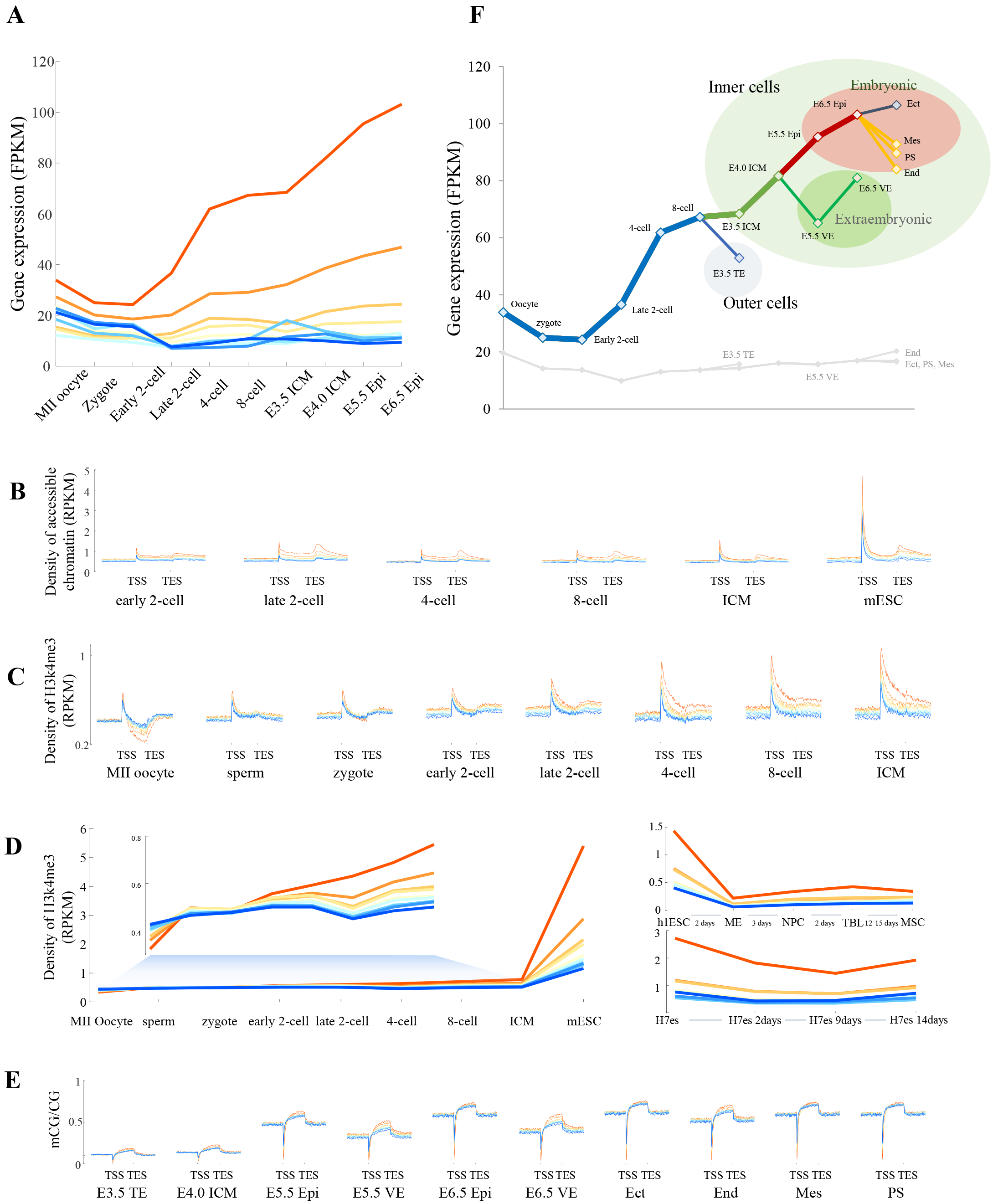
Essential genes in mouse embryos development. Gene expression level for essential genes (red lines) and other groups at each developmental stage. Density of accessible chromatin surrounding TSSs and TESs of genes at each developmental stage. Density of active H3k4me3 modifications surrounding TSSs and TESs of genes at each developmental stage. (D) Dynamics of H3k4me3 density in gene body in preimplantation mouse embryos (left) and postimplantation human embryos (right). (E) CG methylation profiles surrounding TSSs of genes at each developmental stage. (F) Dynamics of gene expression for essential and other genes at each developmental stage. TE, trophectoderm; ICM, inner cell mass; VE, visceral endoderm; Epi, epiblast; Ect, ectoderm; End, endoderm; Mes, mesoderm; PS, primitive streak.

To gain insight into the potential function of essential genes during embryo preimplantation, we examined the gene expression pattern in each developmental stage. Interestingly, differential patterns of transcription were observed during early lineage specification (Fig. 4F). Essential genes were highly expressed in the ICM, which give rise to the entire fetus, while the expression level was much lower in the trophectoderm (TE), the outer layer of the blastocyst stage embryo. During the subsequent formation of primitive endoderm (PE) and epiblast (Epi), essential genes were also higher expressed in embryonic tissues compared to extraembryonic tissues. Finally, during the formation of three germ layers, essential genes were highly expressed in ectoderm, which was derived from the anterior epiblast by embryonic day 6.5 (E6.5), however, essential genes were weak expressed in primitive streak (PE) and PE-derived mesoderm and endoderm (55). Compared to essential genes, nonessential genes were weak expressed and no apparent patterns were observed during embryos development. Thus, transcription of essential genes is indeed required for lineage segregation, especially for the development of fetal-origin part of the placenta.

#### Essential genes are differentially expressed in cancer and normal tissue

Given the fundamental role played by essential genes, it is unsurprising that they represent current and potential novel targets of many antimicrobial and anticancer compounds (56–58). To further investigate the therapeutic implications of essential genes, we examined the relationship between essential genes and cancer genes. We observed a quite variance of cancer genes in different studies (59–63) and more than 2/3 of cancer genes in one study were not cancer genes in another study (fig. S14A). Thus, we compared genes in all these studies, separately, essential genes were significant enriched in cancer genes compared to total protein-coding genes (Fig. 5A). For instance, five famous oncogenes, including *BRCA1*, *BRCA2*, *MYC*, *EZH2* and *SMARCB1*, were essential in human cell survival and associated with chromatin stability, remodeling and modification.

**Figure 5.**
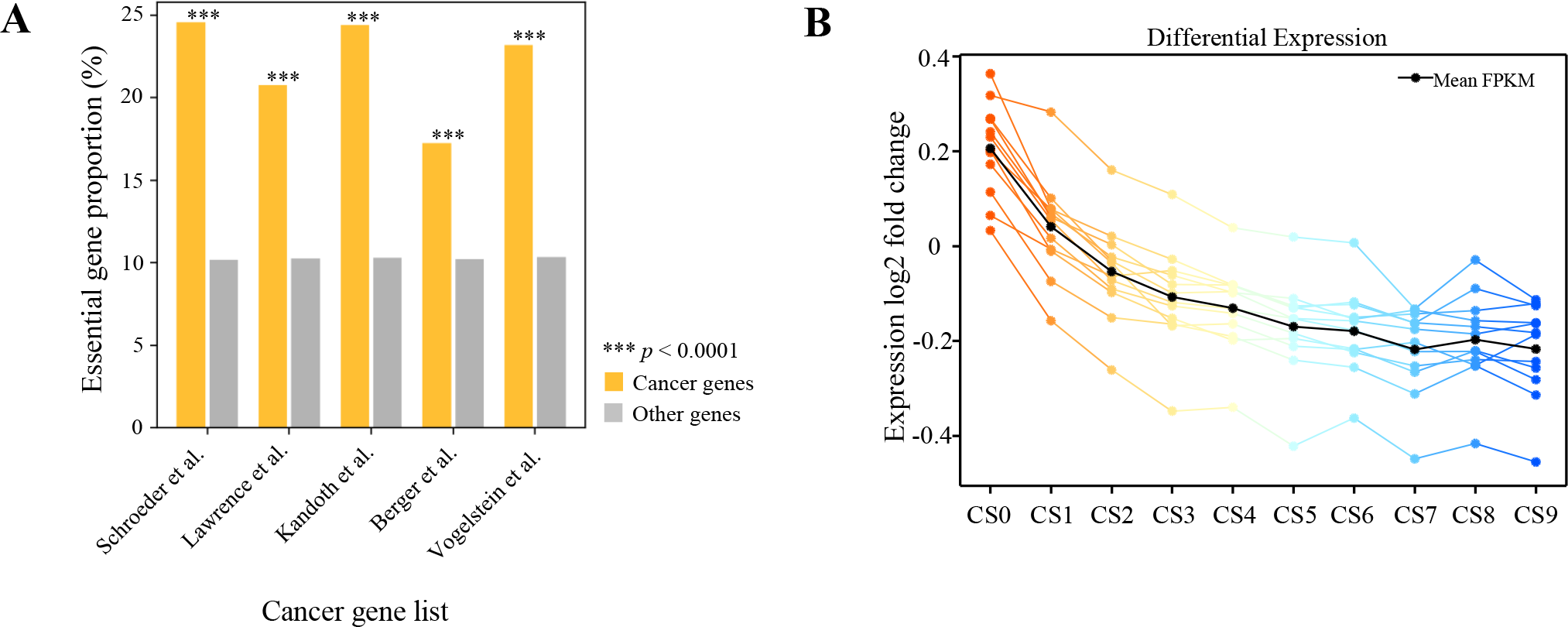
Relationship between human essential genes and cancer. Proportions of human essential genes in 5 cancer gene lists and in all human genes. Significance: *** *P* < 0.0001, fisher exact test). (B) Differential expression of genes in normal and tumor tissues from TCGA database. Each colored line represents mean differential expression fold change (of all donors, from normal to cancer of each donor) of each tumor type.

As reported above, essential genes were higher expressed compared to nonessential genes in both cancer and normal cells (fig. S14B). We next asked whether essential genes were differential expressed between cancer and normal cells. 23 cancer types were examined using gene expression data from TCGA project. Interestingly, the expression level of essential genes were significant higher in cancer cells than that in normal cells (*p*-value = 8.5×10^−6^, paired sample t-test, Fig. 5B). However, nonessential genes exhibit similar transcription activity in both cancer and normal cells. These results suggest that essential genes are more sensitive to tumorigenesis and may further represent superior targets for further drug screening and development.

To further understand the potential function of essential genes in drug screening, we identified 297 significant differential expressed genes (DEGs) using TCGA data (See Methods, Table S1). Using the Drugbank database (64), we then identified 135 candidate drugs for these 297 DEGs (Table S2). Of these candidate drugs, some have already been matured into a drug-targeting strategy for oncology programmes, such as antineoplastic agents: pemetrexed, decitabine, doxorubicin, mitoxantrone and capecitabine. For other candidate drugs, such as anti-infective agents: trifluridine, fleroxacin and antibacterial agents: enoxacin, pefloxacin, ciprofloxacin, may also play a role in cancer treatment and more researches and clinical experiments are required.

### HEGIAP: an interactive web server for studying essential genes

We developed the interactive web server HEGIAP, which integrates abundant analytical and visualization tools to give a multilevel interpretation of gene essentiality for a single gene. HEGIAP provides an overall gene property graph, which shows the gene length, transcript count, and distributions of exons and introns of each transcript. Boxplots are provided for gene properties, including but not limited to gene length, protein length, exon count, and repetitive element count near the promoter region, for the 10 groups of genes (CS0-CS9). The graphs show the corresponding value and group number of each selected gene. For a selected gene, the web server provides histone modification, methylation, and chromatin accessibility profiles and the Hi-C contact map of chromatin structure, all of which have been shown to be significantly correlated with the CS value. Multigene analysis is available to study groups of genes in a comprehensive manner (Fig. 6).

**Figure 6.**
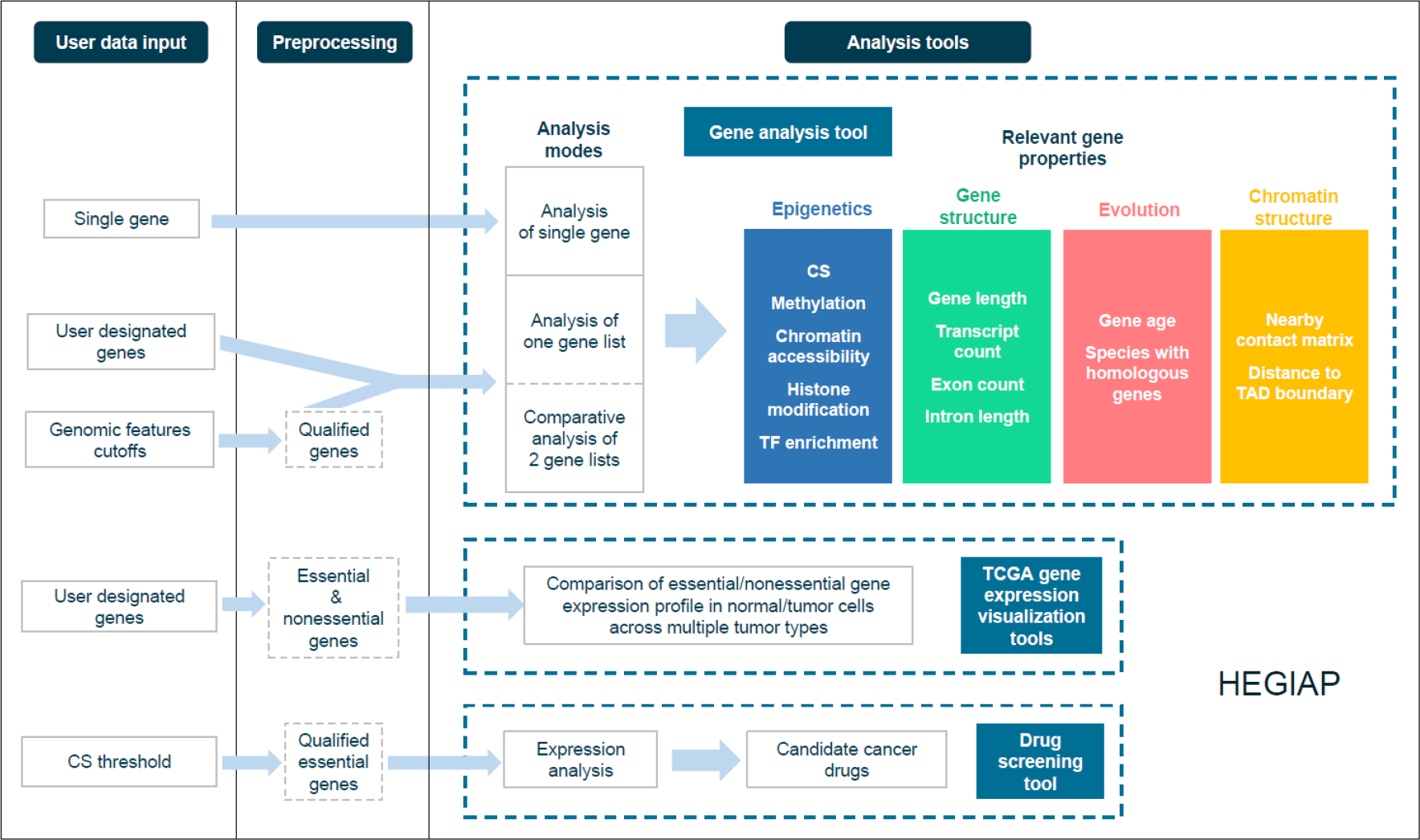
Integrative analysis of individual genes by HEGIAP

HEGIAP supports both feature- and gene-oriented analyses. In feature-oriented analysis, users can obtain all of the genes that meet chosen screening thresholds for multiple features. They can examine the CS distribution or any other property, enabling exploration of possible correlations between CS and other genomic features. In the classic gene-oriented analysis, users specify their chosen genes and are provided a comprehensive view of their essentiality and genomic features. Comparative analysis of two different gene lists is provided to facilitate free exploration of the variation of genomic features between genes of interest or between genes screened by different degrees of essentiality or any other property. Tools are provided to identify genes with aberrant epigenetic modification levels or genomic features based on their essentiality.

As the expression level of essential genes were significant higher in cancer cells than that in normal cells, to facilitate easy examination of this pattern, HEGIAP provide a tool for direct visualization of expression profile across multiple TCGA tumor type of any group of genes uploaded. Genes are also grouped into essential and nonessential sub-groups whose expression profile are also shown for further comparison.

HEGIAP identifies differentially expressed genes in tumor/normal tissues. The web server offers a drug-screening tool, which is based on the assumption that essential genes have predictive power in finding candidate drugs for cancer. Users can set a CS threshold and acquire a list of drugs that significantly suppress expression of cancer-specific highly expressed genes filtered by the CS threshold. The user-friendly interface of HEGIAP was constructed by using R Shiny and requires no plugin installation for users running any popular web browser.

## Discussion

### Three “hubs” location of essential genes

Essential genes have been identified to be important content in multiple life science research including genetic network(65–67)，developmental phenotypes(68), evolution(69), cancer therapy(70), and drug discovery(71). Our work extends the previous studies by revealing a three “hubs” location of essential genes. First, essential genes are “hubs” of PPI networks. As described in the ‘centrality-lethality’ rule that genes and proteins with a high degree of connectivity tend to be essential because their inactivation is more likely to disrupt overall network architecture (48).

Our statistical analysis confirms that gene essentiality is significant correlated with the degree of connectivity and proteins encoded by essential genes are much stable to tolerate translational errors. Second, our work uncovered the structure “hub” of essential genes for the first time. Not only the essential genes are densely clustered in the genome, essential genes are significant clustered in the three-dimensional (3D) structural organization. Third, essential genes are sensitive “hubs” of epigenetic modifications, which also contributes to the high expression of essential genes. As essential genes are centers of both epigenetic medications and chromatin structure, their high transcription activity may further promote the expression of surrounding genes, that is, essential genes may act as the “seeds” of a transcription factory, where endogenous genes are replicated, transcribed, and repaired (72–74). However, more studies are needed to confirm this hypothesis.

### No gene is absolutely essential, but only functions can be so

Consistent with a previous study (42), we confirmed that most essential genes are old genes. However, we also found that an unexpectedly high proportion of the youngest, human-specific genes are essential, and play a role in the regulation of GTPase activity. Although essential genes are highly expressed and genes with high expression level tend to be conserved across species, we noticed a great variance of essential genes in different species. By further examining the PPI networks constructed by essential genes, we found that although gene essentiality changes across species, the biological processes were conserved, this observation provides a new insight into the idea that no gene is absolutely essential, but only functions can be so (48).

### Implications for gene editing and synthetic biology

Major innovations in our ability to edit genome sequences have enabled cost-effective and straightforward genome editing in yeasts, plants and animals (75, 76). The three “hub” locations of essential genes suggest that the effects of gene editing (or gene therapy) on cells should not only consider the target gene and its signaling pathways, but also the associated epigenetic environment and the context of chromatin structure. Further, essential genes can be used as a preferred gene set or important reference of gene interactions for synthetic biology. Additionally, in cancer research, they could facilitate drug discovery in cancer, offer promise as markers in cancer research, and may be useful for identifying clinical therapeutic applications.

In summary, our work provides a very valuable understanding of human essential genes. while due to the limitation of experimental approaches, further work is required not only for understanding the evolutionary plasticity of essential genes across various species, but also for gaining more evidence of the three “hubs” locations of essential genes. These studies will facilitate our understanding of the design principles of transcription regulatory networks, higher-level organization of vital processes and principles underlying drug resistance,

## Materials and Methods

### Dataset

#### For human essential genes

Data on DNase I hypersensitivity (DHS), DNA methylation, histone modifications, CTCF, and evolutionary conservation were downloaded from the ENCODE project and RoadMap database. Hi-C data were obtained from GSE43070 (28) and GSE63525 (29). The TAD boundary were detected according to the protocol described previously (77). Position-specific weight matrices (PWMs) of transcription factors were downloaded from Transfac and Jaspar databases. Data on ncRNAs were downloaded from the NONCODE database (78) (http://www.noncode.org). Essential genes for different species were obtained from DEG (4) (http://www.essentialgene.org/). Cancer data were downloaded from TCGA project. Drug data were obtained from the DrugBank database. Human gene annotations were obtained from the GENCODE database (V21).

#### For essential genes during mouse embryos preimplantation

Assay for transposase-accessible chromatin followed by sequencing (ATAC-seq) were obtained from GSE66390 (79). Histone modification H3K4me3 for mouse early 2-cell, 2-cell, 4-cell, 8-cell stages and ICM were obtained from GSE71434 (80) and the ENCODE project. Histone modification H3K4me3 for H1hESC and H1hESC-derived cells were obtained from previous study (81). Histone modification H3K4me3 for H7es and H7es-derived cells were obtained from the ENCODE project. Mouse gene annotations were obtained from the Mouse Genome Informatics(82).

### Division of genes into groups by CS value

Given the high consistency of essential genes in different cell lines, we used essential genes in KBM7 cell line for this study. Genes were sorted by ascending CS value and divided into 10 groups (CS0-CS9).

### Hi-C data processing

For H1hES cell, Hi-C contact matrices were construct and then normalized using HOMER (http://homer.ucsd.edu/homer/). For GM12878, IMR90 and K562 cell lines, Hi-C contact matrices and loops were obtained from previous study (29) and further normalized using SQRTVC normalization.

### Construction of PPI network

Protein-protein associations were obtained from STRING database (83) (version: 10.5). Using Centiscape (49), we computed specific centrality parameters to describe the network topology and then calculated the connectivity degree for each node in the PPI network. Densely connected regions in large PPI networks were detected using the molecular complex detection method (50). Gene Ontology (GO) analysis was performed using DAVID (84).

### Profiling of epigenetic information

For each gene, genebody as well as 10 kb upstream and downstream were each broken into 50 bins. The ChIP-seq density (RPKM) in these regions was calculated and combined together to get 150 bins spanning 10 kb upstream, the genebody, and 10 kb downstream. The average combined profiles for genes is shown.

### Screening of pan-cancer candidate genes

12 tumor types (COAD, KICH, BLCA, KIRC, CHOL, UCEC, PRAD, KIRP, LIHC, CESC, LUAD, BRCA), from TCGA project were used to screen pan-cancer candidate genes. Differential expression genes (DEGs) were first calculated in each cancer-normal tissue pairs. The 5,000 top scoring DEGs in total genes and the 1,000 top scoring DEGs in essential genes were further combined, and genes in both top scoring DEGs sets were used as candidate genes in each cancer type. Finally, pan-cancer candidate genes were defined as genes that were candidate in at least 8 cancer types.

## Acknowledgments

This work was supported by the Major Research plan of The National Natural Science Foundation of China (No. U1435222), the Program of International S&T Cooperation (No. 2014DFB30020), the National High Technology Research and Development Program of China (No.2015AA020108).

